# Next-generation sequencing based hospital outbreak investigation yields insight into *Klebsiella aerogenes* population structure and determinants of carbapenem resistance and virulence

**DOI:** 10.1101/497958

**Authors:** Adel Malek, Kelly McGlynn, Samantha Taffner, Lynn Fine, Brenda Tesini, Jun Wang, Heba Mostafa, Sharon Petry, Archibald Perkins, Paul Graman, Dwight Hardy, Nicole Pecora

## Abstract

*Klebsiella aerogenes* is a nosocomial pathogen associated with drug resistance and outbreaks in intensive care units. In a 5-month period in 2017, we experienced an increased incidence of cultures for carbapenem-resistant *K. aerogenes* (CR-KA) from an adult cardiothoracic intensive care unit (CICU) involving 15 patients. Phylogenomic analysis following whole-genome sequencing (WGS) identified the outbreak CR-KA isolates to group together as a tight clonal cluster (<7 SNPs apart), suggestive of a protracted intra-ward transmission event. No clonal relationships were identified between the CICU CR-KA strains and additional hospital CR-KA patient isolates from different wards and/or previous years. Genes encoding carbapenemases or drug-resistant plasmids were absent in the outbreak strains, and carbapenem resistance was attributed to mutations impacting AmpD activity and membrane permeability. The CICU outbreak strains harbored an integrative conjugative element (ICE*Kp*10), which has been associated with pathogenicity in hypervirulent *Klebsiella pneumoniae* lineages. Comparative genomics with global *K. aerogenes* genomes showed our outbreak strains to group closely with global ST4 strains, which along with ST93 likely represent dominant *K. aerogenes* lineages associated with human infections. WGS is a powerful tool that goes beyond high-resolution tracking of transmission events into identifying the genetic basis of drug-resistance and virulence, which are not part of conventional diagnostic workflows. With an increasing availability of sequenced genomes from across the globe, population structure analysis offers opportunities to identify emerging trends and dominant clones associated with specific syndromes and geographical locations for poorly characterized pathogens.

## INTRODUCTION

*Klebsiella aerogenes* (formerly described as *Enterobacter aerogenes*) is a ubiquitous member of the *Enterobacteriaceae* family and a significant nosocomial pathogen associated with drug resistance and a wide variety of infections including pneumonia, bacteremia, urinary tract and surgical site infections (1, 2). In vulnerable patients, *K. aerogenes* infections can arise endogenously (gastrointestinal flora) or be acquired from surroundings in the facility where the patient is admitted (horizontal transmission through colonized healthcare workers, contaminated devices/shared equipment, other patients etc.), with the most critical risk factor for acquiring infection being prolonged broad-spectrum antibiotic administration. Additional risk factors for *K. aerogenes* infections include prolonged stay at healthcare facilities especially ICUs and neonatal wards, complex underlying medical conditions, immunosuppression and mechanical ventilation or the presence of foreign devices. Numerous hospital ward outbreaks in both pediatric and adult populations due to *K. aerogenes* have been described due to a common source or spread via patient-to-patient transmission (1, 2). A particularly high frequency of hospital ICU outbreaks was continually reported from Western Europe in a period between the 1990s and early 2000s, that were largely attributed to the spread and endemic establishment of a clonal *K. aerogenes* strain harboring the extended-spectrum B-lactamase TEM-24 (*bla*_TEM-24_) (2).

Within the US and other regions across the globe, *K. aerogenes* has also been reported along with *Klebsiella pneumoniae, Enterobacter cloacae* and *Escherichia coli* to be among the frequently isolated carbapenem resistant *Enterobacteriaceae* (3–5). Clinical CR-KA strains harboring plasmid-borne serine carbapenemases have been described in the US and worldwide, while metallo-β-lactamases and OXA-48 have been reported in Europe, Asia and Brazil (2). However, the primary mechanisms underlying carbapenem non-susceptibility in *K. aerogenes* are thought to be carbapenemase independent and mediated by mutations affecting regulation of chromosomal AmpC β-lactamase expression and membrane permeability (2). The latter has been well documented in *K. aerogenes* with reports describing mutations impacting porin function/expression that can arise *in vivo* during antibiotherapy (6), be reversible (2), and present complex diagnostic and therapeutic management challenges (7).

Despite the role of *K. aerogenes* as an important opportunistic pathogen and its epidemic potential, the clinical relevance of intra-species genetic diversity and significance of specific sequence types (STs) remains unknown. In comparison, in genomically closely related *K. pneumoniae* and to some extent in *E. cloacae*, clonal complexes and sequence types (STs) associated with geographical distribution, multi-drug resistance, hospital outbreaks and disease syndromes have been defined (8, 9). Recently, a multi-locus sequence-typing (MLST) scheme has been developed for *K. aerogenes* (10) that can help explore the above-mentioned issues, however, its performance in evaluating and discriminating clinical/environmental isolates has not yet been reported.

We pursued this study to investigate an outbreak of CR-KA in a cardiothoracic intensive care unit (CICU) at our hospital, which persisted for 5 months despite aggressive infection control measures. The primary goals of our study included whole-genome sequencing (WGS) based investigation of the clonal relationships among the CR-KA strains isolated from patients in our hospital and defining putative loci associated with carbapenem resistance and virulence. In addition, the recently developed publicly available *K. aerogenes* MLST scheme afforded us the opportunity to delineate the population structure of CR-KA stains isolated from patients at our hospital. Our initial findings led us to broadly investigate the origin and significance of specific *K. aerogenes* sequence types identified in our hospital CR-KA strains by performing comparative genomics using publicly available global *K. aerogenes* genomes.

## RESULTS

### Epidemiological and genomic characterization of the CICU CR-KA cluster

In July 2017, five CICU patients had CR-KA isolated from respiratory tract specimens (room occupancy and patient demographics detailed in Fig. 1 and Table S1). The first identified case (Patient A) had a past medical history significant for intravenous drug use and recurrent methicillin-resistant *Staphylococcus aureus* infections. Patient A’s CICU course is described in Fig. S1. The temporal association of subsequent positive cultures in the CICU prompted an outbreak investigation. Between August to November 2017, despite extensive infection prevention interventions, 10 additional CICU patients had positive CR-KA cultures (Table S1). Of the 15 positive cultures from unique patients, 6 were clinical specimens (5 respiratory, 1 blood) and 9 were rectal surveillance cultures. Interventions included contact precautions for all patients on the unit, cohorting of staff, and weekly surveillance cultures of rectal swabs and tracheal aspirates. Environmental surveillance cultures during this period were negative. An epidemiological investigation into the possible risk factors among the CICU patients developing CR-KA infections was non-contributory.

**Fig. 1.**
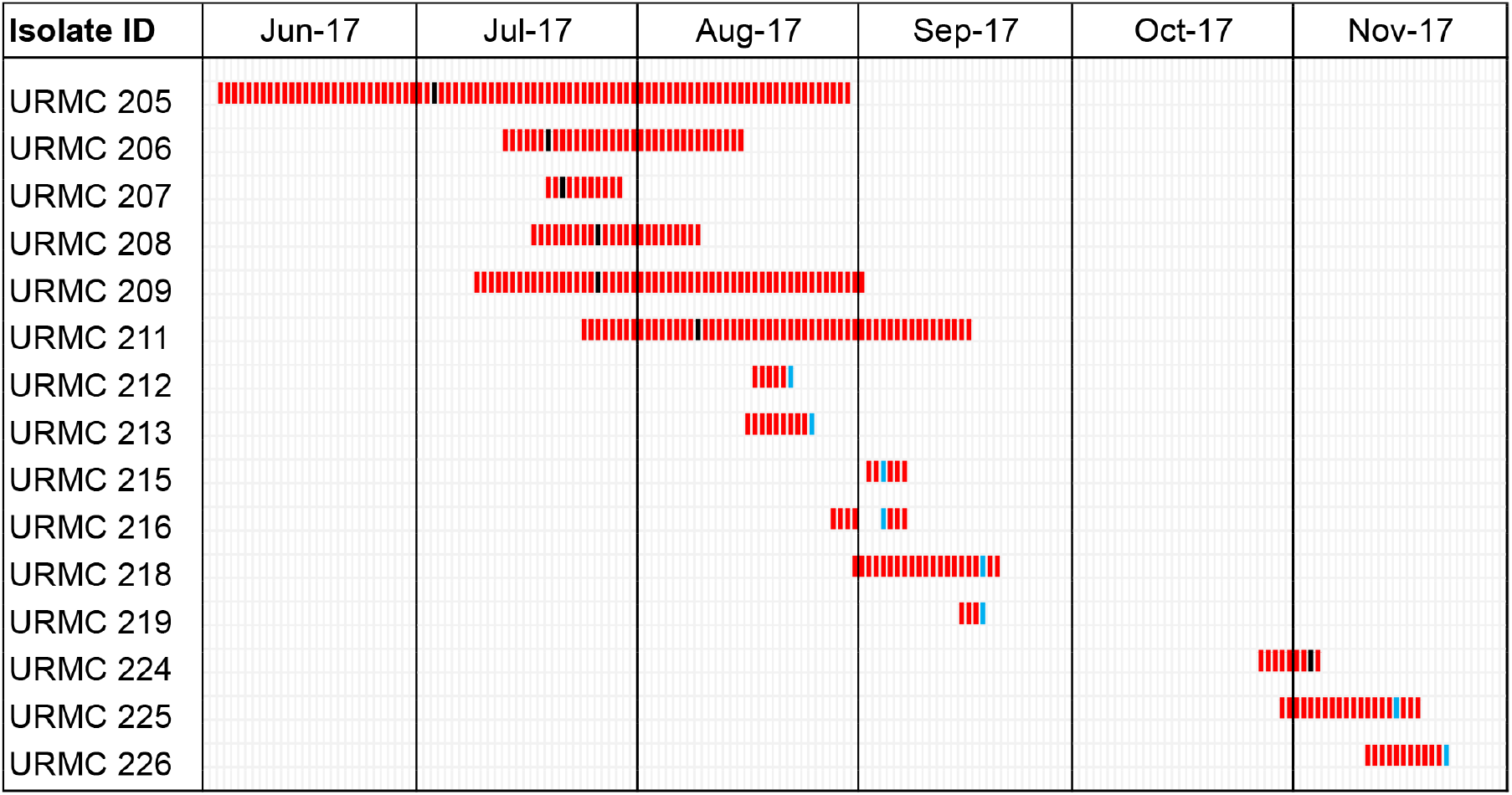
Patient occupancy and overlap in CICU ward during the *K. aerogenes* outbreak. Black bars-first positive CR-KA clinical cultures, blue bars-first positive CR-KA surveillance cultures

Antibiotic susceptibility testing (AST) of the CICU CR-KA strains revealed some differences in phenotype (Table 1) among the carbapenems and cephalosporins. Sixty percent of the CICU cluster strains were resistant to three carbapenems (ertapenem, imipenem and meropenem), while 93% showed phenotypic resistance to ertapenem. Susceptibility to cefepime also varied, with a subset of outbreak strains displaying phenotypic resistance (Table 1).

**Table 1.**
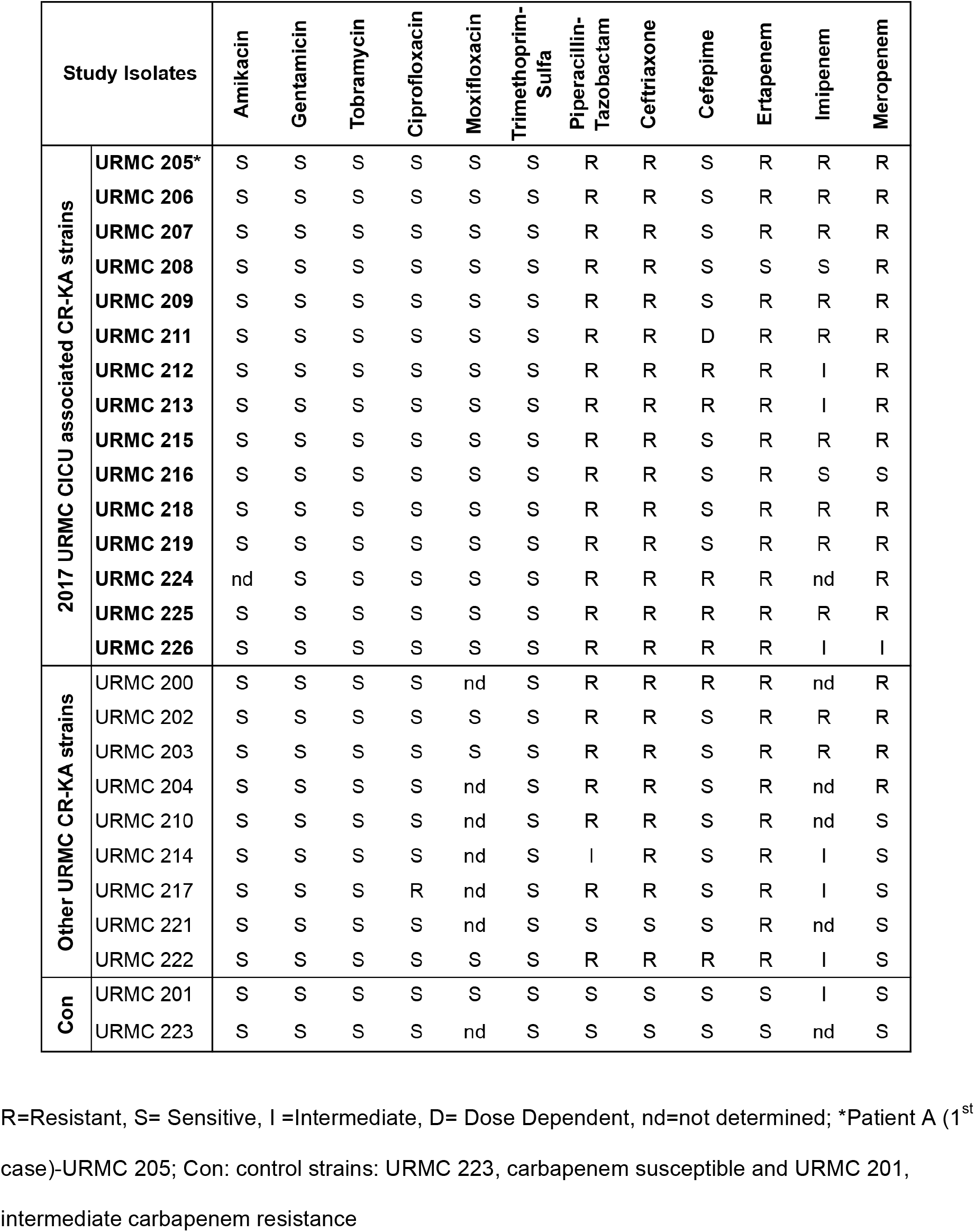
Phenotypic antibiotic susceptibility profiles of *K. aerogenes* strains in this study.

To address the possibility of an outbreak event, particularly in light of variable antibiotic susceptibilities, a WGS investigation was undertaken to identify phylogenetic relationships and transmission links. A total of 26 URMC *K. aerogenes* isolates were sequenced using Illumina WGS. The primary strains investigated in this study were 15 CR-KA isolates from patients admitted in the CICU, between June and Nov 2017 (Table S1). For context and comparison, an additional set of 9 CR-KA strains epidemiologically unlinked to the CICU-cluster strains (patient isolates between years 2015-17), were included in the study (Table S2). Two clinical isolates were also included as controls, URMC 201 (intermediate susceptible to imipenem) and URMC 223 (susceptible to all carbapenems tested). The AST profiles of the non-CICU CR-KA study strains are described in Table 1.

All of the 26 sequenced URMC *K. aerogenes* genomes showed high coverage (>88%) relative to the *K. aerogenes* reference KCTC 2190 strain (ATCC 13048^T^), Dataset 1. Single nucleotide polymorphisms (SNPs) were identified across the study genomes relative to the reference sequence (pair-wise SNP differences ranged from 1-28,170, see Dataset 2). MLST assignment indicated that all of the CICU clinical and surveillance isolates belonged to ST4, and SNP-based analyses grouped them in a tight cluster separately and distantly from non-outbreak isolates (Fig. 2, Dataset 2). Within the CICU cluster, strains differed from URMC 205 (first case, patient A isolate) by less than 7 SNPs. In addition, these isolates bore an identical plasmid profile (based on replicon and plasmid typing) (Table S3). In contrast, all of the 2017 non-CICU isolates were significantly distant from the CICU outbreak isolates (>20,000 SNPs). The most closely related non-CICU CR-KA strain was URMC 201 (isolated in 2015). This strain was also ST4, with 433 SNPs in a pairwise comparison to URMC 205. (Fig. 2, Dataset 2). These relationships, coupled with the epidemiological data, indicated that the 2017 CICU CR-KA strains were of clonal origin.

**Fig. 2.**
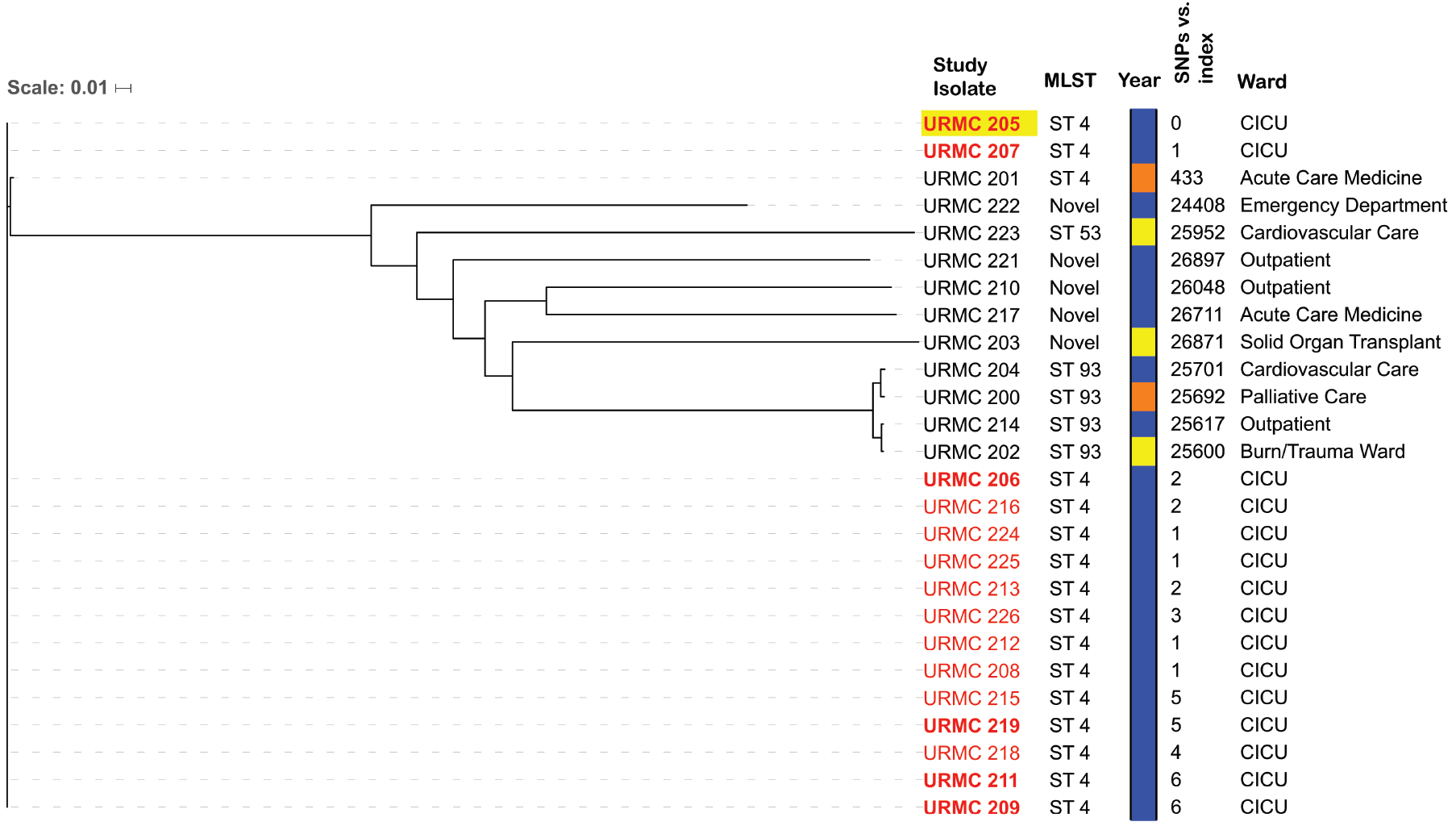
Dendogram showing pairwise SNP differences based phylogenetic relatedness of URMC *K. aerogenes* strains. Whole genome sequence of *K. aerogenes* KCTC 2190 (ATCC 13048) was used for reference mapping. A total of 28,170 discriminatory high-quality SNPs in the core genomes obtained by the CFSAN SNP pipeline were used to plot the tree (excluding mobile elements and putative recombination sites). CICU outbreak strains: IDs highlighted in red, clinical isolates in bold, patient A isolate (1^st^ case) with yellow background. Year strain isolated: orange-2015, yellow-2016, blue-2017. SNP differences relative to patient A shown. Scale bar indicates nucleotide substitutions per site.

### Carbapenem resistance in the URMC CR-KA strains was driven by adaptive chromosomal gene alterations

#### A. WGS based identification of acquired antibiotic resistance genes

Despite phenotypic carbapenem resistance, the CICU CR-KA did not harbor genes for carbapenemases, extended-spectrum β-lactamases, or plasmid-borne AmpC cephalosporinases. A single non-outbreak CR-KA isolate (URMC 203) harbored a carbapenemase gene (*bla*_nmcA_). No horizontally acquired genes conferring resistance to non-β-lactam antibiotics were identified among the study strains, consistent with their susceptibility profiles (Table 1).

#### B. Non-synonymous sequence alterations in key chromosomal loci implicated in carbapenem resistance

In the absence of genes encoding carbapenemases and ESBLs in the CR-KA outbreak strains, variations in other genetic loci associated with carbapenemase-independent resistance mechanisms were investigated (11–13). Focusing on the AmpC cephalosporinase and outer membrane porins, sequence variations in *ampD, ampG, ampR, omp35, omp36* and *ompR* genes in the study strains were assessed relative to the ‘wild-type’ allele in the carbapenem-susceptible reference type strain KCTC 2190 (14). The sequences were also compared to alleles in URMC 223 (carbapenem susceptible) and URMC 201 (intermediate susceptibility). For the *omp* genes, the upstream DNA sequences were also assessed. The *ampG* and *ompR* genes were wild-type in all study strains. Variants were identified in all other loci and are described below:

***ampD***. Mutations were identified in the *ampD* gene for each of the 24 CR-KA isolates in this study (Table 2), while the control strains URMC 223 and URMC 201 bore the ‘wild-type’ *ampD* allele. The outbreak strains harbored single missense SNPs (either 284G>T or 482G>A) in *ampD* resulting in Trp95Leu or Arg161His substitutions. Six non-outbreak CR-KA strains harbored independent non-synonymous single substitutions while 2 missense substitutions were identified in URMC 221. A single non-outbreak strain, URMC 202, harbored a nonsense mutation resulting in a truncated AmpD protein (Table 2). DNA sequence corresponding to *ampD* allele was absent in isolate URMC 203, (the only study strain to possess a carbapenemase-encoding gene *bla*_NMC-A_) due to a large deletion in the genomic region harboring *ampD* and the neighboring *ampE* gene. The potential impact of Trp95Leu or Arg161 His substitutions on AmpD activity in the outbreak strains was investigated by homology-based structural modeling of *K. aerogenes* AmpD using a high-resolution crystal structure of *Citrobacter freundii* AmpD (15), a close homolog (83.33% amino acid sequence identity). AmpD contains a hydrophobic surface to accommodate its GlcNAc-anh-MurNAc ligand, which is made up of tripeptide and glycan moieties. The tripeptide portion of GlcNAc-anh-MurNAc is coordinated through three salt bridges between carboxyl groups on the tripeptide and residues Arg71, Arg161 and Arg107, while the peptide backbone is oriented across the hydrophobic surface. Trp95 forms a planar surface at the end of the ligand-binding channel to position the diaminopimelate moiety at the distal end of the tripeptide. Based on the *in silico K. aerogenes* AmpD model, the positively charged guanidinium group of Arg161 forms two strong electrostatic interactions with the carboxyl group of D-glutamine on the tripeptide portion of the ligand (Fig. 3). Mutation of this residue to histidine was predicted to weaken ligand binding, likely affecting the positioning of the ligand in the active site. In the Trp95Leu mutation, the hydrophobicity in the region is preserved, but the shorter length of the leucine side chain leaves a gap at the end of the binding channel likely affecting the positioning of the entire ligand (Fig. 3). These observations lend support that the missense mutations observed in the CICU outbreak strains would alter ligand docking on AmpD likely reducing/inhibiting its activity. Four out of the seven CR-KA non-outbreak strains had substitutions within glycan and peptide interacting regions of AmpD (Table 2), suggesting that these alterations might also impact activity.
***ampR***. All of the outbreak strains harbored the reference *ampR* allele. Several non-outbreak CR-KA strains harbored 2-5 substitutions that likely represent variant alleles as they were also observed in the control carbapenem-susceptible strain URMC 223. A single non-outbreak CR-KA strain, URMC 210, bore a nonsense mutation in the *ampR* gene resulting in a premature stop codon (Trp117X), likely resulting in a non-functional truncated AmpR protein.
***omp35*** and ***omp36***. Among the 15 outbreak CR-KA strains, three different *omp36* variants were identified relative to the wild-type allele (Table 3). An identical profile of missense SNPs in these strains and the control strain, URMC 201 (intermediate carbapenem susceptibility) were observed relative to the reference genome allele. Seven outbreak strains had additional mutations that resulted in severely truncated proteins. Several non-outbreak CR-KA strains also harbored missense SNPS of unclear significance. Two strains URMC 202 and URMC 204 harbored distinct frame-shift mutations yielding truncated Omp36 protein variants. A 42 base-pair region of high variation (nucleotides 680-724), corresponding to 15/16 amino acid substitutions in loop L5 was observed in all the clinical isolates relative to the reference sequence (Table 3, Fig. S2). The hypervariable region results in a different charge profile in the region and has been previously reported in a *K. aerogenes* study describing imipenem resistant clinical isolates (harboring ESBL TEM-24) from patients in France (16). The predicted Omp36 protein in clinical strain URMC 221 and the carbapenem susceptible control strain (URMC 223) bore 87% identity relative to the reference genome Omp36 and were considered significantly distant variants (not included in the comparative analyses). All but one of our 24 CR-KA strains had the wild-type allele of *omp35*. A single strain (URMC 202) bore a deletion in the N-terminal encoding region. The DNA sequence upstream of the *omp35* gene was investigated to identify mutations in the promoter sites of the strains, of which one (URMC 204) had a nucleotide difference of unclear significance at the -22 position.

**Table 2.** Non-synonymous SNPs in *ampD* gene associated with carbapenem resistance in the URMC *K. aerogenes* isolates.

| Study Isolates |  | Differences in *ampD* gene relative to wild-type allele |  | Effect on encoded AmpD sequence |  |
| --- | --- | --- | --- | --- | --- |
|  |  | MS SNPs | NS SNPs | Single AA substitutions | PS |
| **CICU outbreak associated CR-KA strains** | URMC 205*, URM C 207, URM C 209, URM C 211, URM C 212, URM C 213, URM C 215, URM C 218, URM C 219, URM C 224, URM C 224, URM C 226 | 482G>A |  | Arg161His |  |
|  | URMC 206, URM C 208, URM C 216 | 284G>T |  | Trp95Leu |  |
| **Outbreak unrelated CR-KA strains** | URMC 200 | 338T>G |  | Ile113Ser |  |
|  | URMC 202 |  | 412C>T |  | Gln138X |
|  | URMC 204 | 117C>T |  | Pro39Ser |  |
|  | URMC 210 | 335C>T |  | Ser112Leu |  |
|  | URMC 214 | 280G>A |  | Ala94Thr |  |
|  | URMC 217 | 496G>C |  | Gly166Arg |  |
|  | URMC 221 | 478A>G, 501C>A |  | Ile160Val, Ala168Asp |  |
|  | URMC 222 | 492T>G |  | Glu164Asp |  |
| **Control strains** | URMC 201, URM C 223 |  |  |  |  |

No sequencing reads mapped to the *ampD* gene in URM C 203.
\*Index patient isolate.
Abbr: SNP: single nucleotide polymorphism, MS: missense, NS: nonsense, AA: amino acids, PS: premature stop codon

**Table 3.** Non-synonymous genetic lesions in *omp36* genes and resulting alterations in Omp36 sequence in the URMC *K. aerogenes* isolates.

| Study Isolates |  | Differences in *omp36* gene sequence relative to wild-type allele |  |  |  | Effect on encoded Omp36 amino acid sequence relative to wild-type sequence |  |  |  |  |
| --- | --- | --- | --- | --- | --- | --- | --- | --- | --- | --- |
|  |  | FS | HV region (Nt 680-724) | NS SNPs | MS SNPs | FS | HV region (AA 226-241) | PS | Truncated | Single AA substitutions in protein synthesized |
| **CICU outbreak associated CR-KA strains** | URMC 205*, URMC 207 | + | + |  | 175A>G, 564T>C, 566A>G | pAsp91ThrfsX12 |  | 102X | + | Ile59Val |
|  | URMC 206, URMC 208, URMC 212, URMC 213, URMC 216, URMC 224, URMC 225, URMC 226 |  | + |  | 175A>G, 564T>C, 566A>G |  | + |  |  | Ile59Val, Asp189Gly |
|  | URMC 209, URMC 211, URMC 215, URMC 218, URMC 219 |  | + | 184A>T | 175A>G, 564T>C, 566A>G |  |  | 62X | + | Ile59Val |
| **Outbreak unrelated CR-KA strains** | URMC 200, URMC 203, URMC 210, URMC 214 |  | + |  | 564T>C, 566A>G, 615 T>G, 834T>C, 835A>G |  | + |  |  | Asp189Gly, Asp205Glu, Asn279Asp |
|  | URMC 202 | + | + |  | 564T>C, 566A>G, 615 T>G, 834T>C, 835A>G | pTrp77LysfsX2 |  | 78X | + | Trp77Lys |
|  | URMC 204 | + | + |  | 564T>C, 566A>G, 615 T>G, 834T>C, 835A>G | pSer304ProfsX21 | + | 324X | + | Asp189Gly, Asp205Glu, Asn279Asp |
|  | URMC 217 |  | + |  | 564T>C, 566A>G, 569T>A |  | + |  |  | Asp189Gly, Phe190Tyr |
|  | URMC 222 |  | + |  | 175A>G, 564T>C, 566A>G, 615 T>G |  | + |  |  | Ile59Val, Asp189Gly, Asp205Glu |
| **Control strain** | URMC 201 (intermediate carbapenem R) |  | + |  | 175A>G, 564T>C, 566A>G |  | + |  |  | Ile59Val, Asp189Gly |

\*Index patient isolate.
Abbr: SNP: single nucleotide polymorphism, NS: nonsense, MS: missense, FS: frame-shift, PS: premature stop codon, HV: hypervariable region, Nt: nucleotides, AA: amino acids

**Fig. 3.**
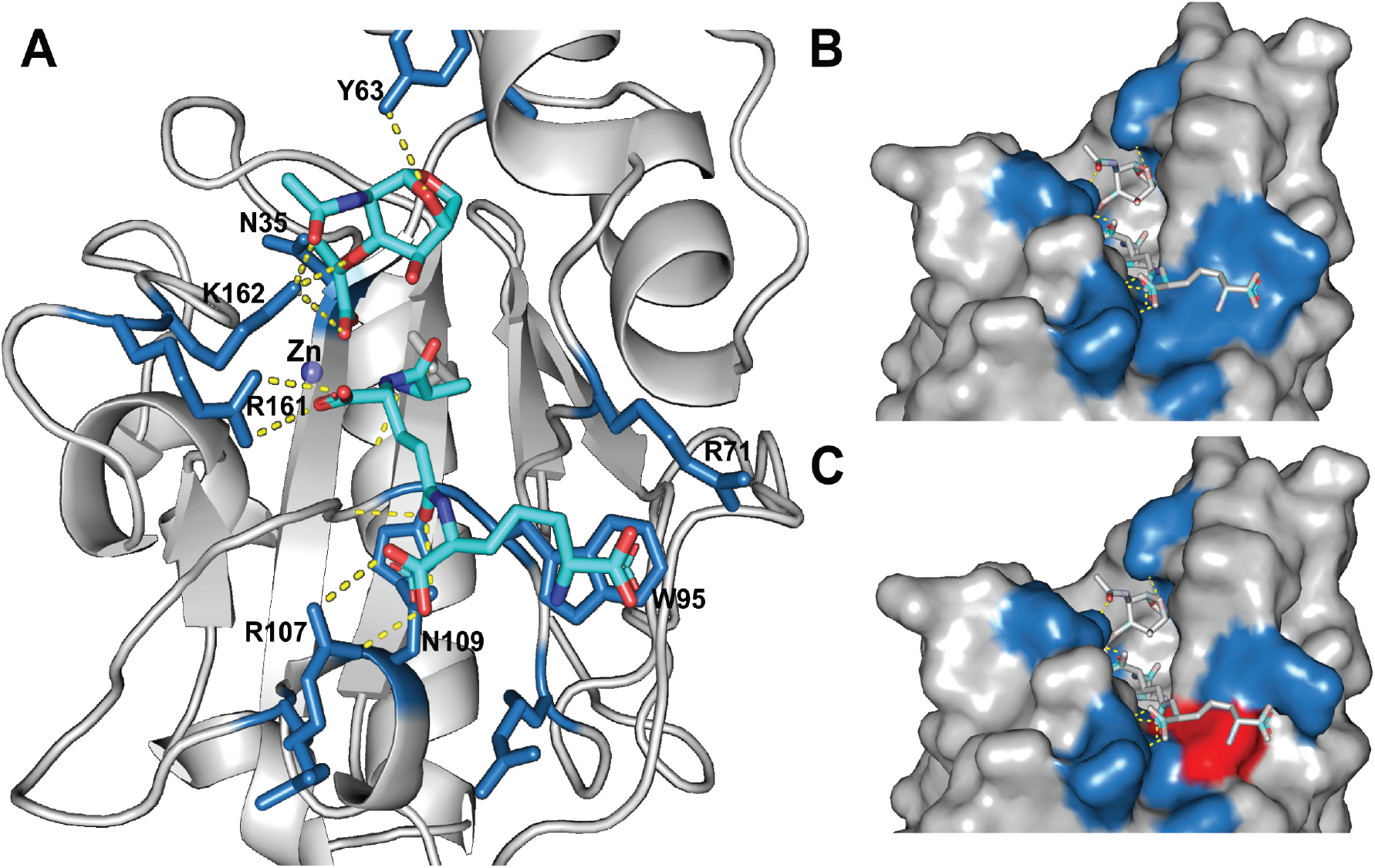
Computational modeling of the impact of *ampD* mutations on AmpD in URMC outbreak CR-KA strains. A) *K. aerogenes* AmpD modeled on *C. freundii* AmpD structure. Key residues interacting with glycan and peptide portions of ligand are shown. B, C) Surface models of *K. aerogenes* AmpD depicting the wild-type binding surface for the diaminopimelate moiety (W95, top) versus the binding surface of AmpD containing the W95L mutation (blue surfaces indicate amino acid positions from panel A; bottom, altered surface highlighted in red).

### Virulome analysis identifies a large pathogenicity-associated integrative and conjugative element (ICE), ICE*Kp*10 in the CICU outbreak CR-KA strains

The prolonged nature of the clonal CR-KA outbreak in our CICU led us to search for putative virulence genes or pathogenicity loci, which could have promoted persistence and transmission. A cluster of chromosomally encoded genes encoding yersiniabactin (*ybt*) metallophore and colibactin (*clb*) genotoxin systems was identified in all of the outbreak strains (Fig. 4). These loci have been implicated in invasive infections of *K. pneumoniae* (17).

**Fig. 4.**
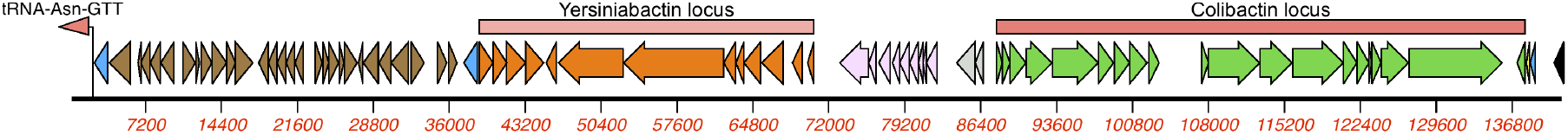
Yersiniabactin and colibactin encoding gene loci on integrative conjugative element ICE*Kp*10 in the CICU outbreak CR-KA strains. Blue arrows integrase encoding genes, brown arrows-Zn^2+/^Mn^2+^ modules grey arrows-genes encoding mobilization proteins, pink arrows: vir-T4SS system.

The *ybt* locus harbored putative genes involved in regulation as well as synthesis of the siderophore, corresponding transport associated proteins, and a receptor protein for the uptake of metal-bound siderophore. The *clb* locus included putative homologs encoding enzymes, transferases and transport proteins involved in production and secretion of the polyketide colibactin (Fig. 4, Dataset 3). Investigation of the genomic loci associated with the virulence factor gene cluster identified them to be present on a mobilizable integrative conjugative element (ICE) inserted in a tRNA-Asn site adjacent to a gene encoding the glycine cleavage system (Fig. 4). The ICE element bore a modular arrangement of gene clusters encoding mobile elements, P4 like integrase, type IV secretion system conjugation machinery and mobilization genes (Fig. 4, Dataset 3). The element was identified to be ICE*Kp*10, using a recently described typing and interpretation scheme of *ybt* and *clb* sequences (17).

Among the URMC non-outbreak *K. aerogenes* strains, these putative virulence genes were identified in only 4/10 isolates, which were either ST4 or ST93. The control strain URMC 223 harbored the yersiniabactin locus exclusively. Detailed descriptions of these elements in the URMC *K. aerogenes* strains are described in Table S4.

### Comparative analyses of URMC CR-KA and publicly available *K. aerogenes* genomes

To gain insights into the emergence and epidemiology of the CICU outbreak clones and to place our hospital CR-KA strains in the broader context of global *K. aerogenes* strains, comparative phylogenomic analyses were performed using Harvest genomics suite (18). Publicly available *K. aerogenes* genome assemblies (n=110) were included in the analyses. These included 70 clinical and surveillance strains isolated from human specimens, 3 ‘environmental’ strains, and 36 strains of unknown origin (Dataset 4). Based on the newly described MLST scheme, ST4 and ST93 strains were found to be markedly overrepresented in the available genomes (51.8%, 57/110).

Excellent correlation was observed between Harvest generated tree topologies (Fig. 5) as well as pairwise SNP differences (Dataset 5) as compared to those generated by the CFSAN SNP pipeline, for the URMC CR-KA study strains. Based on Harvest analyses, the CICU outbreak strains clustered closely with each other and to other global ST4 genomes, as compared to the other URMC CR-KA strains (outbreak-unrelated), which were distantly dispersed throughout the phylogenomic distribution (Fig. 5). The MLST based sequence types of URMC CR-KA and the global *K. aerogenes* genomes also correlated tightly with HARVEST generated core-genome based topologies (Fig. 5).

**Fig. 5.**
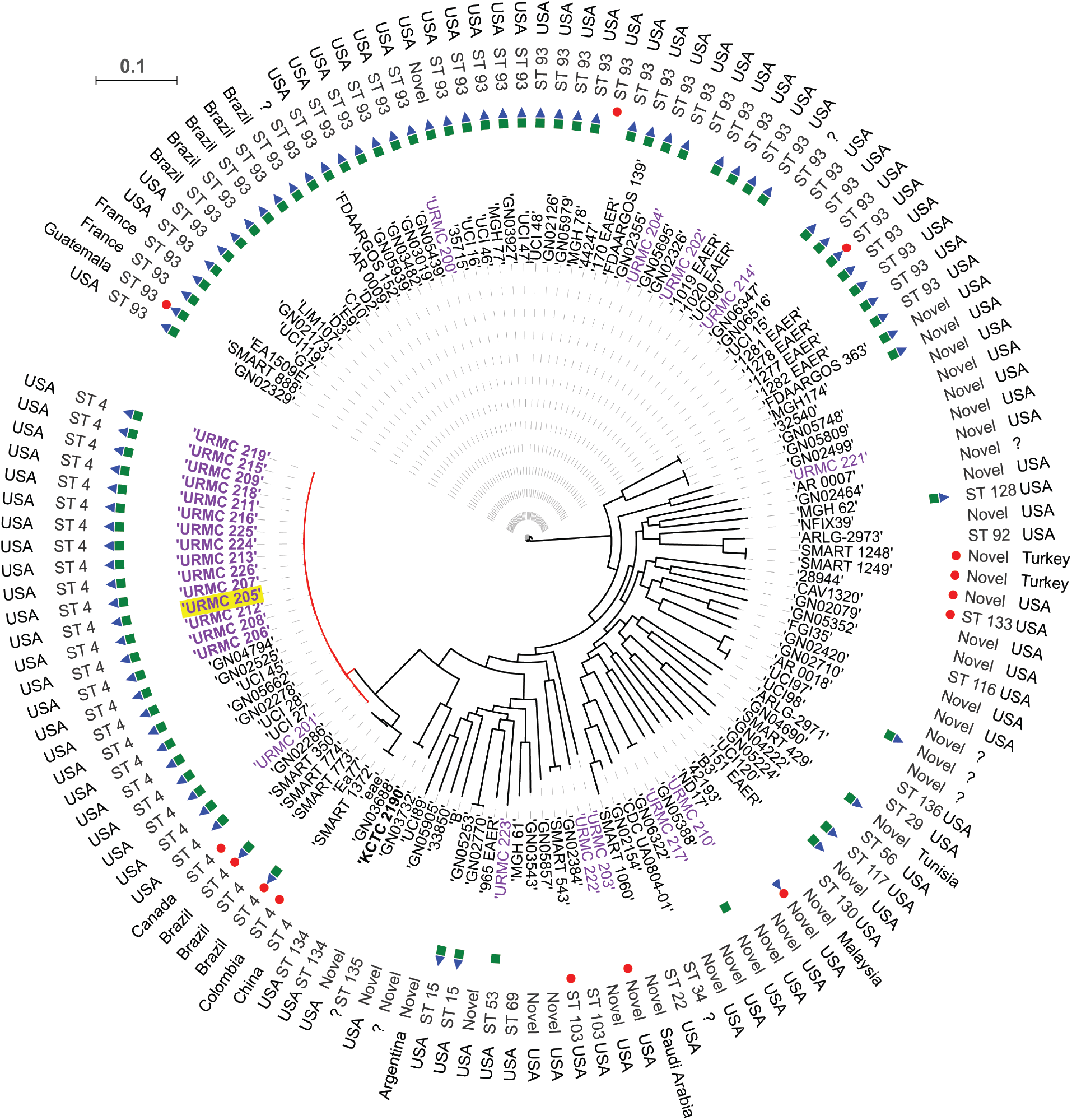
Harvest based phylogenomic comparisons of URMC *K. aerogenes* genomes with global *K. aerogenes* genomes. Discriminatory SNPs based on core genome comparisons were used to plot the tree. URMC study *K. aerogenes* strains in purple (outbreak strains in bold, index patient strain highlighted yellow). Presence of genes encoding yersiniabactin siderophore system (green squares), colibactin synthesis cluster (blue triangles), and carbapenamases (red circles) in assembled genomes shown. Scale bar indicates nucleotide substitutions per site.

Six publicly available assembled genomes grouped closely with the CICU outbreak strain genomes (< 200 SNPs apart). These included ST4 strains, UCI 27, UCI 28, UCI 45, which have been described in a carbapenem resistance surveillance study by Cerqueria et al. (19), and GN04794, GN05662 and GN02525; strains derived from varied US patients’ clinical specimens (blood, sputum, wound drainage) Fig. 5, Dataset 4). The above-mentioned strains had fewer SNP differences in relation to the CICU outbreak strains as compared to URMC 201, the closest and only non-outbreak ST4 strain isolated in our hospital (Fig. 5, Dataset 5).

Carbapenemase encoding genes were identified in a total 13/111 global *K. aerogenes* genome assemblies (11.8%) (Fig. 5). These included genes encoding KPC-2 (n=7), KPC-3 (n=1), OXA-48 (n=4) and NDM-6 (n=1). The chromosomal serine carbapenemase, *bla*_NMC-A_, was solely present in URMC 203, a CR-KA strain isolated from a patient in our hospital in 2015. Among the thirty ST4 genomes, 4 strains (13%) harbored genes encoding carbapenemases (4/4, KPC-2), and these were clinical strains isolated from non-USA patients. The relative contribution of carbapenemase mediated versus non-carbapenemase mediated mechanisms of resistance to carbapenems in the global *K. aerogenes* could not be assessed due to the absence of antibiotic susceptibility metadata for most strains.

A characteristic genomic feature of the URMC CICU outbreak strains and a subset of non-outbreak associated strains was the presence of the yersiniabactin siderophore and colibactin systems. Using Kleborate (17), we investigated the distribution and organization of these systems in the global *K. aerogenes* genomes to identify associations, if any, with specific ST types and geographical regions (Fig. 5, Datasets 6). The prevalence of yersiniabactin and colibactin encoding systems in the global *K. aerogenes* genomes was found to be 53.64% (59/110) and 52.72% (58/111) respectively. A 100 % association was found between the presence of colibactin in the genomes and concurrent presence of yersiniabactin. Higher prevalence of the virulence cluster was observed in ST4 and ST93 types; 85% (12/14) and 95% (42/44) respectively, although these two ST types were also the most abundantly represented in the available set of genomes. The other ST types were less well represented in the study set (n<3), so the prevalence of these systems in them could not be accurately established. Interestingly, the strains exclusively designated ‘environmental’ isolates (B3, FGI35, and B) did not harbor genes encoding the above-mentioned virulence systems (Fig. 5, Dataset 6). These trends correlated with analyses of our 26 hospital strains, where 75% of the ST 93 strains (3/4) were positive for yersiniabactin and colibactin systems, while none of the strains with novel/unassigned ST types harbored genes encoding the same (0%, 5/5).

## DISCUSSION

Whole genome sequencing presents a powerful resource that can be deployed for prospective and comprehensive outbreak investigations offering tremendous resolution in tracking transmission events and delineating genomic determinants associated with drug resistance and virulence (20). This WGS study was initiated in order to establish the molecular epidemiology of CR-KA strains isolated from patients in our hospital. Beyond the outbreak investigation, whole genome data enabled us to address additional fundamental questions associated with carbapenem resistance, virulence attributes and population structure of *K. aerogenes* that are poorly understood.

Fifteen patients were associated with the CICU outbreak event in the period between July-November 2017. Based on the differences in AST patterns in a subset of study strains, it was initially questioned whether the strains were clonal. WGS investigations demonstrated that the 15 CICU CR-KA isolates differed from each other by less than 7 SNPs and grouped distantly from the other hospital CR-KA isolates, which were used a baseline for context and comparison (Fig. 2). Based on these findings, the CICU CR-KA strains were concluded to be part of a single clonal cluster indicating protracted intra-ward transmission. Environmental sampling within the CICU remained negative and a common source of the outbreak or contributory risk factors could not be identified. After the last positive patient surveillance culture (Nov 2017), no CR-KA strains were isolated in a 2-month period of continued surveillance and the outbreak was deemed to have subsided. Phylogenomic analysis (albeit limited) of the non-CICU CR-KA isolates did not identify specific dominant clones circulating in the hospital (Fig. 2, Dataset 2).

Infections due to carbapenem-resistant organisms present complex diagnostic and therapeutic management challenges and a better understanding of how adaptive or acquired resistance emerges in the healthcare environment is needed (21). In clinical CR-KA strains, carbapenem resistance has been associated with carbapenemase production or adaptive mutations following antibiotic exposure (2). Among our 24 study CR-KA isolates, a single non-CICU strain harbored a carbapenemase-encoding gene, *bla*_NMC-A_. NmcA has been reported in *E. cloacae* (22), but to our knowledge this is the first report of this carbapenemase being identified in *K. aerogenes*. All but one of the 24 CR-KA strains were found to harbor mutations in genes involved in the synthesis or regulation of the inducible AmpC cephalosporinase (*ampD, ampR*) and outer membrane porins (*omp 35, omp36*). A majority of mutations were found to be within the open-reading frames of *ampD* and *omp36* (catalogued in Tables 2 and 3).

There are limited reports regarding the role of AmpD in adaptive carbapenem resistance in *K. aerogenes* strains (23, 24), although the role of AmpD in AmpC expression in *E. cloacae* has been well established (25). *In silico* modeling of *K. aerogenes* AmpD using the *C. freundii* AmpD crystal structure (15) predicted that Arg161 His and Trp95Leu substitutions due to SNPs in the outbreak strains would likely impact enzymatic activity (Fig. 3). Other *ampD* mutations in several non-CICU CR-KA isolates were found to result in substitutions that could affect substrate-enzyme interactions (Table 2).

Mutations in *omp36* have been described in clinical CR-KA strains and functional studies investigating their impact have been reported (6, 26). A diverse array of mutations was identified in the *omp36* gene among the CR-KA study strains, including non-synonymous mutations resulting in frame-shifts or premature stop codons resulting in truncated and likely non-functional Omp36 variants (Table 3). Additional SNPs relative to the carbapenem-susceptible reference genome that resulted in substitutions of unknown significance in predicted β-sheets and extracellular loop regions (Table 3, Fig. S2).

Even within the CICU CR-KA clonal cluster, heterogeneity was identified in *ampD* and *omp36* genes (Table 2, Table 3). These genetic loci likely represent mutational hotspots associated with adaptive carbapenem resistance in *K. aerogenes*. It is likely that the CICU CR-KA cluster represents a population of clonal origin, albeit with micro-heterogeneity in the above regions. The testing and archiving of single isolated carbapenem resistant colonies instead of multiples during individual patient specimen workup likely represents a limitation that did not allow us to capture the full complement of CR-KA strain microdiversity associated with individual patients during the outbreak event.

The detailed significance of the alleles and mutations described above in development of carbapenem resistance in *K. aerogenes* needs to be verified by additional genetic (allelic exchange and complementation) and biochemical approaches. Moreover, complex regulatory networks including transcriptional activators, sensor kinases and two-component systems have been implicated in adaptive drug resistance in *Enterobacteriaceae spp*. (13), and their roles in contributing towards carbapenem resistance in our study strains cannot be ruled out.

Despite their lack of horizontally acquired drug resistance elements, the outbreak strains in this study managed to persist for several months in the face of an active infection prevention effort prompting us to assess their virulence determinants. The outbreak strains and a subset of non-outbreak CR-KA strains harbored an ICE encoding the metallophore yersiniabactin (Ybt) and genotoxin colibactin (Clb) systems (Fig. 4, Table S4). A Ybt encoding pathogenicity island has been described in association with a prolonged nation-wide outbreak in the Netherlands involving multiple hospitals and >100 patients due to a multi-drug resistant *Enterobacter hormaechei* clone (27). A recent study by Lam *et al*. described the prevalence of *ybt* to be particularly high (>80%) in certain hypervirulent *K. pneumoniae* clonal-groups (CG23) (17). The study also reported a significant association of Ybt with an increased risk of invasive infections (bacteremia, liver abscesses etc.). Ybt was first described in pathogenic *Yersinia spp*., encoded by a chromosomal gene cluster termed the High Pathogenicity Island (HPI), with a critical role in iron scavenging during infection (28). Subsequent studies reported acquisition of the HPI in other clinical *Enterobacteriaceae* strains (17, 29) and additional functions ascribed to Ybt include evasion of host lipocalin-2 (30) and sequestration/import of heavy metals (31). The polyketide colibactin is frequently associated with yersiniabactin and has been shown to induce chromosomal instability and DNA damage in eukaryotic cells (32). Our findings present new avenues for research investigating the role of ICE elements encoding Ybt and Clb in the virulence of clinical *K. aerogenes* strains. While horizontal transmission of carbapenemases via mobile elements is increasingly recognized as a major public health issue (33), the transmission of mobilizable virulence factors present an underappreciated threat in the healthcare environment that warrants more surveillance.

In order to set our hospital strains (outbreak and non-outbreak) into a broader context, core-genome comparisons and MLST were used to examine the population structure of our strains relative to global *K. aerogenes* strains pulled from public databases (Fig. 5, Datasets 5, 6). This analysis is the first evaluation of the nascent *K. aerogenes* MLST scheme in discriminating clinical *K. aerogenes* isolates. The scheme was found to be robust with distribution of STs correlating closely with topologies based on *K. aerogenes* strains core-genomes. Our preliminary analyses suggest that ST4 and ST93 might be dominant global clones associated with *K. aerogenes* infections. Outbreak CR-KA strains clustered closest to other clinical US ST4 strains, suggesting a clonal expansion of this ST (Fig. 5). It is noteworthy that the ST4 group also included carbapenemase-producing *K. aerogenes* isolates from international sites. These included CR-KA strains associated with drug-resistant intra-abdominal and urinary tract infections in patient samples from Brazil, Canada, Colombia and China that had been sequenced as a part of large surveillance study, SMART (Study for Monitoring Antimicrobial Resistance Trends) (34). ST93 was the most prevalent sequence type in the global *K. aerogenes* assembled genomes (43/110, 39%), with a wide geographical distribution including the US (Fig. 5). Four of the eleven non-CICU CR-KA strains from our hospital belonged to this group. Two closely related ST93 isolates, *K. aerogenes* 1509E and G7, have been described as representatives of clonal strains associated with multiple multi-drug resistant *K. aerogenes* outbreaks in France (35, 36).

Incidentally, global ST4 and ST93 isolates also had a higher prevalence of HPI encoding *ybt* and *clb* (85% and 95% respectively). The success of these potentially “high-risk” clones needs to be examined more closely by undertaking large-scale studies with strains from diverse global sites and patient populations. These studies will help examine niche adaptation, emergence of antibiotic resistance and evolution of pathogenicity leading to a better understanding of *K. aerogenes*.

In summary, genomic approaches for surveillance and outbreak investigations are emerging as critical functions for infection prevention and diagnostic microbiology laboratories. Apart from evaluating the effectiveness of infection measures and dimension of transmission events, WGS applied across sets of isolates is a powerful tool for assessing virulence factors and identifying clinically relevant and emerging sequence types.

## MATERIALS AND METHODS

### Setting, study design, *K. aerogenes* strains and metadata

The University of Rochester Medical Center (URMC) is an 830-bed tertiary-care medical center, with a 14-bed cardiothoracic intensive care unit, serving the Greater Rochester Area, New York. Following approval by the University of Rochester Institutional Review Board (RSRB00068143), a total of 26 *K. aerogenes* strains isolated from patients at URMC in the course of regular clinical care and/or surveillance efforts were selected for the study. Each isolate corresponded to a single first CR-KA strain isolated during the course of hospitalization. For context and comparison, an additional set of *K. aerogenes* strains epidemiologically unlinked to the CICU outbreak was included in the study. Ward occupancy and pertinent clinical and epidemiological information were obtained through review of patient medical records and the laboratory information system, and are described in Tables S1 and S2.

### Antibiotic susceptibility testing (AST) of the study *K. aerogenes* strains

AST of the study strains was performed as part of routine diagnostics using the VITEK^®^2 (BioMérieux, France) system and/or Kirby Bauer disk-diffusion methods. AST interpretations were based on interpretive criteria defined by the Clinical and Laboratory Standards Institute (M100 document, 27^th^ Edition).

### Sequencing library preparation and raw data acquisition

The isolates were cultured on standard laboratory media from archived frozen stocks, examined for purity, and re-identified by Vitek MALDI-TOF MS (BioMérieux, France). Single colony genomic DNA extractions were performed using MagNA Pure Compact instrument (Roche Diagnostics, Indianapolis, IN); the DNA quality was analyzed using QuantiFluor dsDNA system (Promega). Nextera XT kit (Illumina, San Diego, CA) was used for preparing dual-indexed WGS libraries. For QC dsDNA quality of individual samples was analyzed using the 4200 TapeStation system (Agilent). Following library cleanup, individual libraries were pooled at equimolar ratios and denatured; DNA concentration was determined using Qubit ssDNA kit (Thermo fisher Scientific). The libraries were combined with20pM PhiX control and sequenced on the Illumina Miseq™ benchtop sequencer (Illumina, San Diego, CA) at the URMC Genomic Core Facilities, using V3 kit and 2X300 bp paired-end protocol.

### Genomic analyses

Analyses were performed using an in-house bioinformatics pipeline ‘URMC Bacterial Genomic Analysis Pipeline’ (v2.0.6), run on a high-performance computer cluster at the Center for Integrated Research Computing at University of Rochester. **A) Quality control:** For each sample, the read quality scores across all bases were assessed using FastQC v0.11.5 (37) and low quality reads were trimmed using Trimmomatic v0.36 (38). Genomic coverage of the sequencing reads relative to the reference *K. aerogenes* KCTC 2190 genome (ATCC 13048^T^; Refseq accession number: NC_015663.1) were determined using Bowtie 2 v2.2.9 (39). Across the study isolates, alignments showed average genomic coverage of 90.84% (minimum 89%) with an average depth of coverage: 100X (excluding regions below 12X). Quality metrics are detailed in Dataset 1. **B) Core-genome SNP calling workflow:** A read mapping approach was used to assess SNPs in the genomes of the study CR-KA strains relative to the reference genome *K. aerogenes* KCTC 2190. Mapping, variant calling and phylogenetic analysis were performed by locally installed CFSAN SNP Pipeline v1.0.0 (40), an analysis workflow developed by the U.S Food and Drug Administration (FDA). CFSAN pipeline employs a 2-phase variant calling workflow and the ‘optimized’ version of the pipeline with criteria as applied by Saltykova et al (higher stringency in allele frequency thresholds and coverage) (41) was applied. In the first phase, variants were called based on *mpileup function* of SAMTools and *mpileup2snp tool* from VarScan (minimum average base quality=20, minimum read depth of coverage at site=12, minimum allele frequency=90%). Densely clustered SNPs that could arise due to recombination were excluded (>/=3 SNPs in a 50 bp window). High-confidence SNP variants meeting the criteria were composed to a list. In the 2^nd^ phase, nucleotide sites at the listed positions were determined for all sites (minimum allele frequency threshold for SNP filtering=90%). SNPs located in mobile elements as annotated by RAST were excluded. The final nucleotide sites post filtering corresponding to the listed positions were used to build a SNP matrix. A multi FASTA file with concatenated SNP matrix entries were used for inferring phylogeny by FastTree (42), which were visualized using ITOL (43). **C) Sequence assembly, scaffolding, annotation and analyses:** De novo genome assemblies (average N_50_: 344943 across all isolates) were generated by SPAdes genome Assembler v3.11.1 (44), and the assembly quality was assessed by QUAST v4.5 (45). Ordering and orientation of contigs was performed using Medusa. Draft genomes were annotated with RAST (46) and Prokka (47). MLST on the *K. aerogenes* genomes was performed using a newly developed publicly available scheme (10). Alleles and genetic markers corresponding to acquired antibiotic resistance, virulence and plasmid replicons were identified using PlasmidFinder (48), ResFinder (49), and Virulence Factor Database (50). Typing of genetic loci associated with mobilizable yersiniabactin siderophore and genotoxin colibactin systems was performed using Kleborate (17), and the loci were visualized with MacVector software (MacVector, Cary, NC). Sequence variations in chromosomal genes associated with carbapenem resistance in the study strains were assessed relative to alleles in control strains (URMC 201, URMC 223) and ‘wild-type’ alleles in carbapenem susceptible (14) reference genome KCTC 2190 [Genbank ID-*ampD*: 10792472 (EAE_11350), *ampG*: 10792757 (EAE_12735), *ampR:* 10792632 (EAE_12115), *omp35*. 10793271 (EAE_15245), *omp36:* 10795060 (EAE_24205) and *ompR:* 10791222 (EAE_05245)]. Multiple sequence alignments of gene alleles and corresponding proteins was performed using Vector NTI software (Invitrogen, Carlsbad, CA) and the alignments were manually inspected to identify substitutions that would likely impact function. **D) Comparative genomics of publicly available global ***K. aerogenes*** assembled genomes with URMC CR-KA assemblies:** Using a custom shell script, publicly available *K. aerogenes* genomes available as of July 2018 in the NCBI genome database were downloaded from the NCBI FTP site. Harvest genomics suite was used to perform intraspecific core-genome alignments as described before (18); phylogenies were visualized using ITOL (43). **E) Accession number(s):** WGS and metadata corresponding to the study URMC *K. aerogenes* isolates were deposited at NCBI under BioProject accession number PRJNA504784. Accession numbers for individual isolates are listed in Table S1.

### *In silico* protein analyses

For homology modeling, SWISS-MODEL (51) was used to thread *K. aerogenes* AmpD (acc WP_015704411.1, aa1-187) through the structure of *C. freundii* AmpD (15) (PDB 2y2c, aa1-187). The overall quaternary structure of *K. aerogenes* AmpD was predicted with high precision (95% confidence, 99% coverage). Comparative analyses and imaging of protein structures were performed with PyMOL (52). JalView was used to create alignments (53).

## ACKNOWLEDGEMENTS

We are grateful to the URMC Clinical Microbiology Laboratories and Infection Prevention staff in specimen processing, data collection and epidemiological investigations. We wish to acknowledge the URMC Genomics Research Center for support with WGS. We also thank Steve Gill (URMC, Genomics Research Center) for reviewing the manuscript draft. Internal funding from University of Rochester Department of Pathology and Laboratory Medicine supported this study. We report no conflicts of interest relevant to this article.

## Supplementary Data

**Fig. S1.**
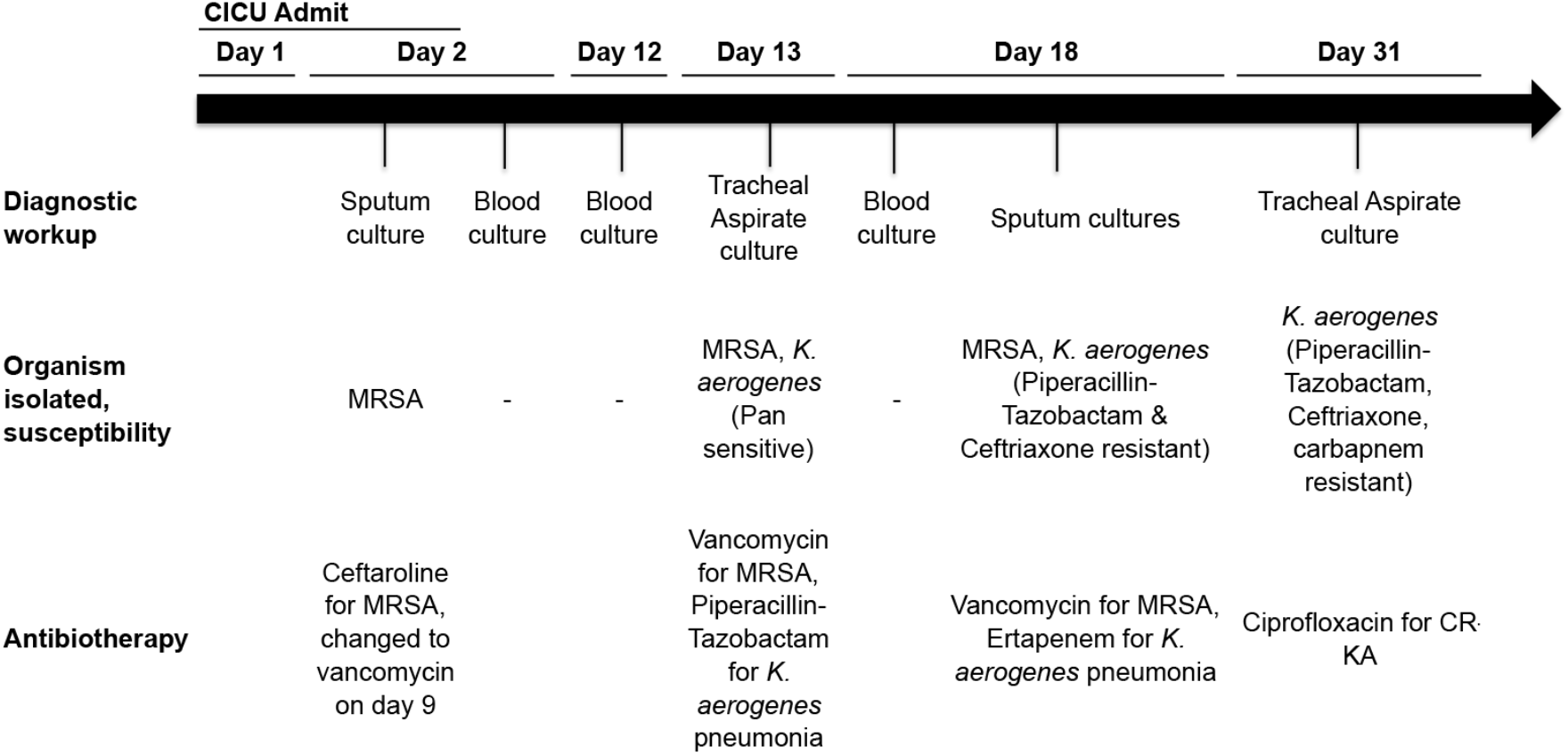
Patient A timeline following CICU admit highlighting diagnostic workup, microbiological findings and antibiotic administration prior to development of CR-KA respiratory infection.

**Fig. S2.**
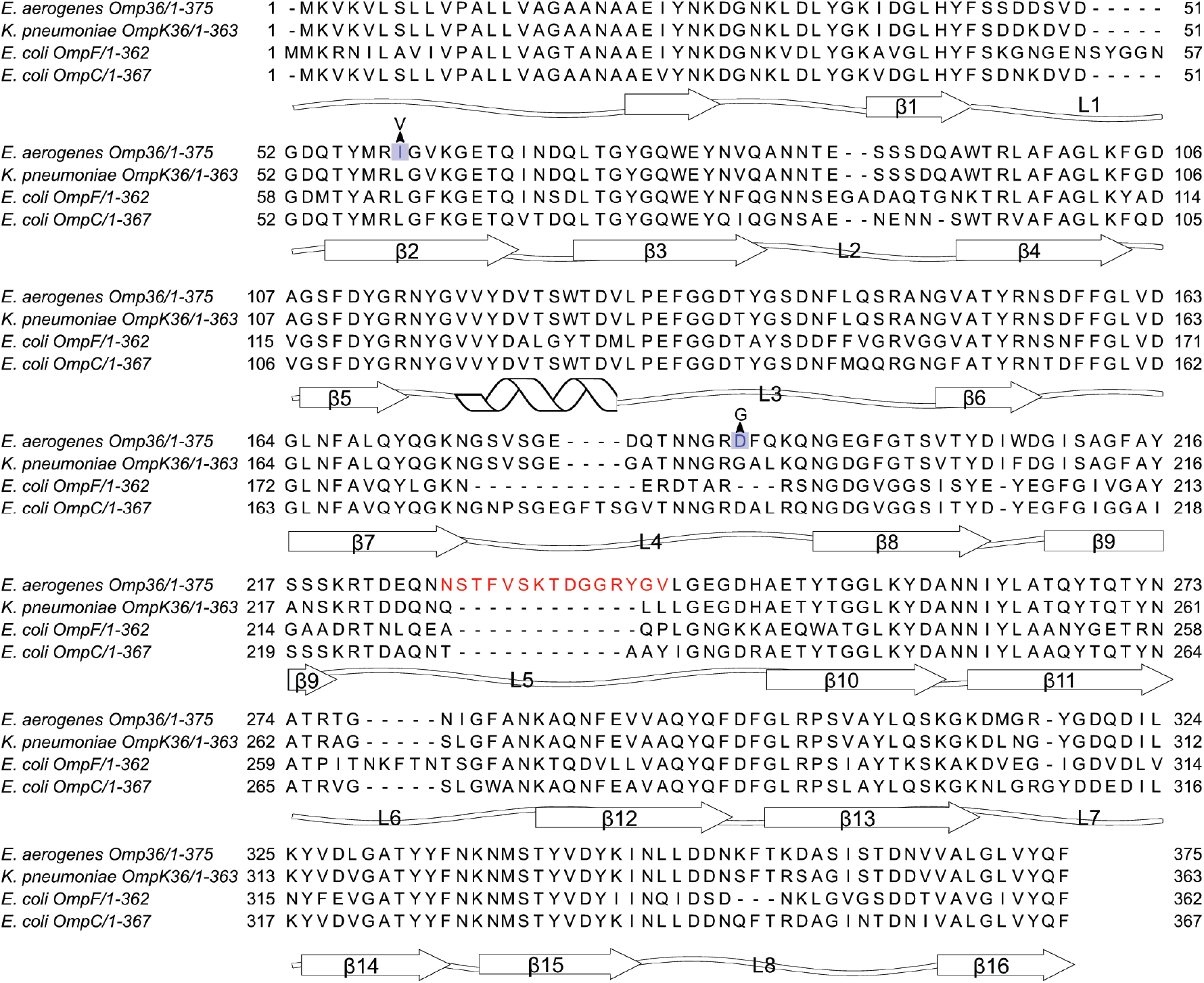
Comparison of *K. aerogenes* Omp36 with representative porins of the OmpC and OmpF families. Predicted β-sheets, loops and helixes are shown. Non-synonymous SNP mutations in CICU outbreak CR-KA strains resulting in substitutions relative to reference highlighted in blue.

**Table S1:**
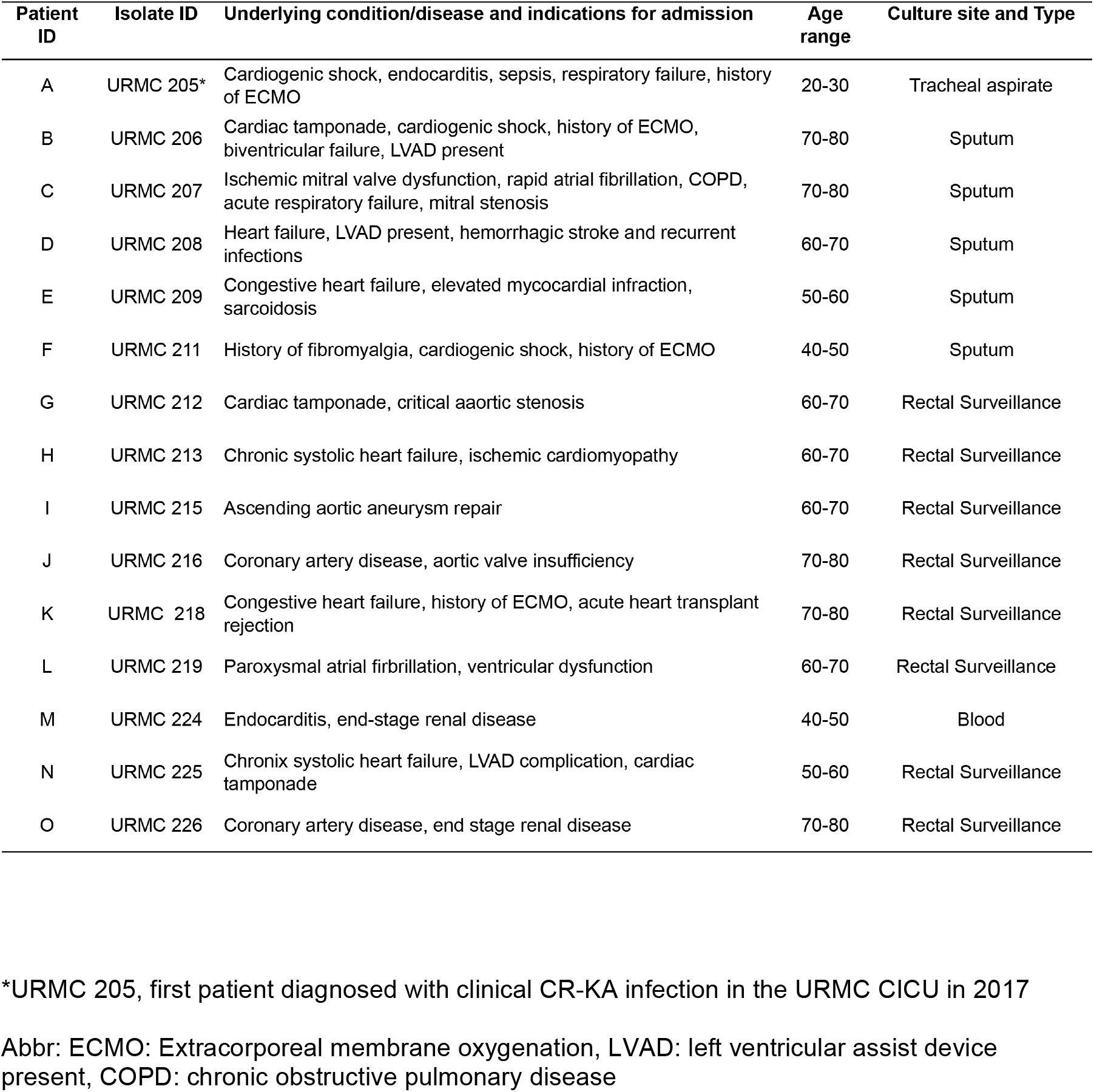
Attributes of patients associated with the 2017 URMC CICU CR-KA cluster.

**Table S2:** Summary of additional hospital *K. aerogenes* strains in this study. (isolates not associated with CICU CR-KA cluster)

| Isolate ID | Year of positive culture | Gender | Age range | Culture site and type | Patient's unit/ward |
| --- | --- | --- | --- | --- | --- |
| URMC 200 | 2015 | M | 20-30 | Catheter urine | General Medicine |
| URMC 201 | 2015 | M | 30-40 | Blood | Acute Care Medicine |
| URMC 202 | 2016 | M | 60-70 | Broncholaveolar lavage | Burn/Trauma |
| URMC 203 | 2016 | M | 70-80 | Biliary drain fluid | Solid Organ Transplant |
| URMC 204 | 2016 | M | 70-80 | Catheter urine | Cardiovascular Progressive Care |
| URMC 210 | 2017 | F | 80-90 | Urine | Outpatient |
| URMC 214 | 2017 | F | 90-100 | Urine | Outpatient |
| URMC 217 | 2017 | F | 70-80 | Urine, cytосcopy | Acute Medicine |
| URMC 221 | 2017 | F | 90-100 | Urine | Outpatient |
| URMC 222 | 2017 | F | 40-50 | Abdominal ascites fluid | Emergency Department |
| URMC 223* | 2017 | F | 70-80 | Urine | Cardiovascular Step Down |

\* All strains except URMС 223 and URMС 201 were deemed carbapenem resistant by CDC and CLSI definitions
URMC 223 was a pan-sensitive *K. aerogenes* isolated from the urine of a patient in the CICU in Oct 2017 (within the outbreak period)

**Table S3:**
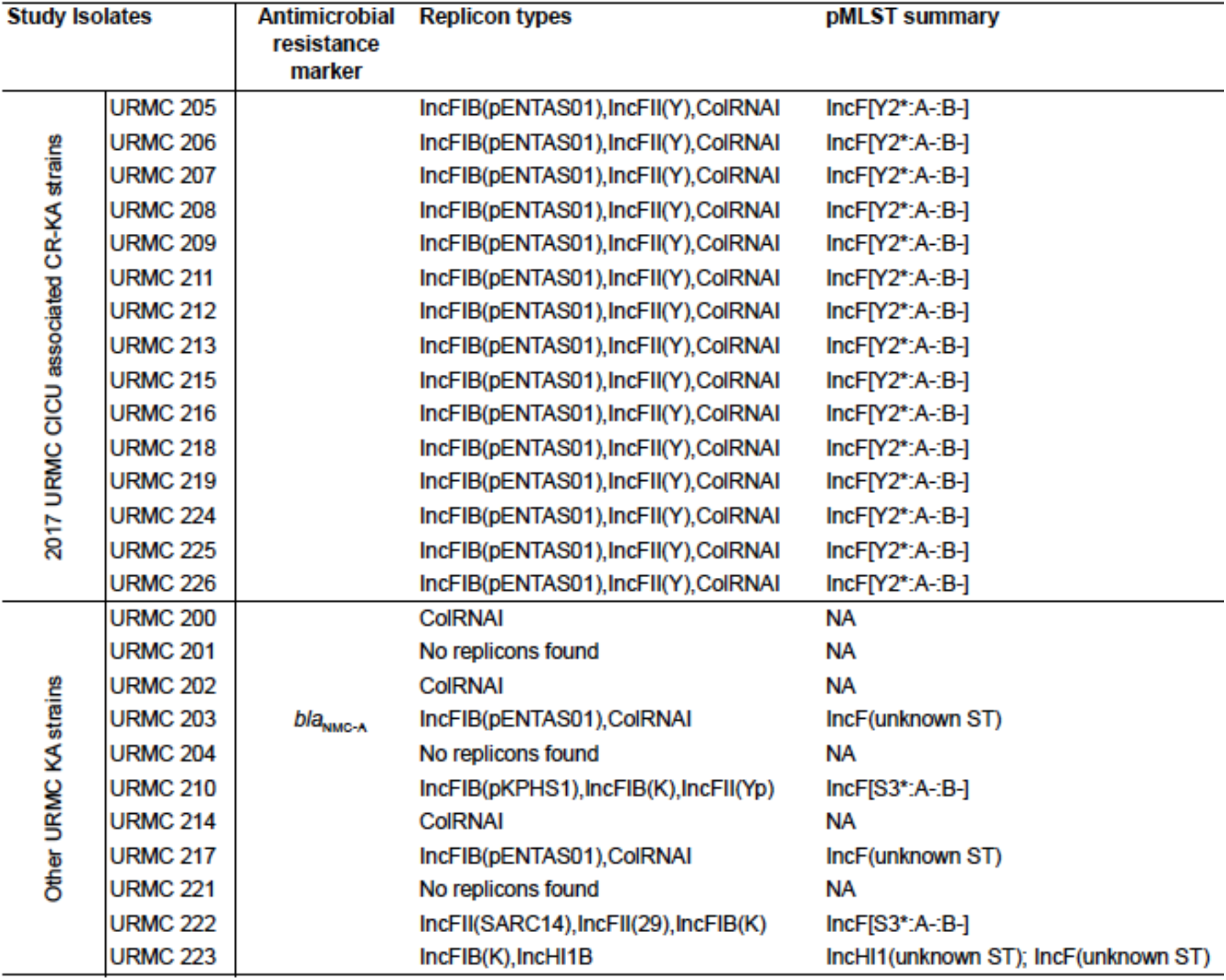
Acquired antibiotic resistance markers, plasmid replicon types and plasmid multilocus sequence typing of the study *K. aerogenes* strains.

**Table S4:** Typing yersiniabactin and colibactin encoding gene clusters in URMC *K. aerogenes* strains using Kleborate.

| Study Isolates |  | Yersiniabactin | YbST | Colibactin | CbST |
| --- | --- | --- | --- | --- | --- |
| 2017 URM C ICU associated CR-KA strains | URMC 205 | ybt 17; ICEKp10 | 289-1LV | clb 3 | 17-3LV |
|  | URMC 206 | ybt 17; ICEKp10 | 289-1LV | clb 3 | 17-3LV |
|  | URMC 207 | ybt 17; ICEKp10 | 289-1LV | clb 3 | 17-3LV |
|  | URMC 208 | ybt 17; ICEKp10 | 289-1LV | clb 3 | 17-3LV |
|  | URMC 209 | ybt 17; ICEKp10 | 289-1LV | clb 3 | 17-3LV |
|  | URMC 211 | ybt 17; ICEKp10 | 289-1LV | clb 3 | 17-3LV |
|  | URMC 212 | ybt 17; ICEKp10 | 289-1LV | clb 3 | 17-3LV |
|  | URMC 213 | ybt 17; ICEKp10 | 289-1LV | clb 3 | 17-3LV |
|  | URMC 215 | ybt 17; ICEKp10 | 289-1LV | clb 3 | 17-3LV |
|  | URMC 216 | ybt 17; ICEKp10 | 289-1LV | clb 3 | 17-3LV |
|  | URMC 218 | ybt 17; ICEKp10 | 289-1LV | clb 3 | 17-3LV |
|  | URMC 219 | ybt 17; ICEKp10 | 289-1LV | clb 3 | 17-3LV |
|  | URMC 224 | ybt 17; ICEKp10 | 289-1LV | clb 3 | 17-3LV |
|  | URMC 225 | ybt 17; ICEKp10 | 289-1LV | clb 3 | 17-3LV |
|  | URMC 226 | ybt 17; ICEKp10 | 289-1LV | clb 3 | 17-3LV |
| Other URM C KA strains | URMC 200 | ybt 0; ICEKp10 | 220-2LV | clb 3 | 13-2LV |
|  | URMC 201 | ybt 17; ICEKp10 | 289-1LV | clb 3 | 17-1LV |
|  | URMC 202 | ybt 17; ICEKp10 | 267 | clb 3 | 19-1LV |
|  | URMC 203 | - | 0 | - | 0 |
|  | URMC 204 | ybt 16; ICEKp12 | 277-3LV | clb 3 | 17-1LV |
|  | URMC 210 | - | 0 | - | 0 |
|  | URMC 214 | - | 0 | - | 0 |
|  | URMC 217 | - | 0 | - | 0 |
|  | URMC 221 | - | 0 | - | 0 |
|  | URMC 222 | - | 0 | - | 0 |
|  | URMC 223 | ybt 4; plasmid | 221-3LV | - | 0 |

Abbr: YbST: Yersiniabactin locus sequence type, CbST: Colibactin locus sequence type

